# Learned landmark associations support online visual control under degraded visibility

**DOI:** 10.64898/2026.07.13.738310

**Authors:** Grace C. Roessling, Brett R. Fajen

## Abstract

Humans and other animals often act in environments that are at least partly familiar, where aspects of the spatial layout are known. Although such knowledge is known to support navigation and spatial cognition, its role in the online control of action remains unclear. We investigated whether drivers use knowledge of road layout to guide steering in high and low visibility and, if so, the form of such knowledge. In two simulated driving experiments (total N = 90), participants repeatedly drove winding roads containing segments with and without fog. Drivers who repeatedly experienced the same road exhibited more stable steering and lane positioning than drivers encountering novel roads, but only when visibility was reduced. These advantages were accompanied by superior performance on post-tests assessing knowledge of road geometry. We next examined the form of such knowledge by dissociating global knowledge of road layout from local associations between landmarks and road segments. Disrupting landmark-road segment associations produced the largest impairment in steering performance. The benefits of prior experience were largely preserved when road-segment order was scrambled but landmark associations remained intact. These findings show that spatial knowledge can support moment-to-moment steering control when visibility is reduced. Rather than relying on a globally coherent representation of the environment, drivers use local associations between landmarks and upcoming road geometry to anticipate future demands. More broadly, the results elucidate how familiarity with environmental structure contributes to the control of action when visual information is degraded, revealing a close interplay between spatial knowledge and visual control.

**Significance Statement:** People routinely act within surroundings they have encountered before, from commuting on the same streets to walking familiar hallways. Whether the spatial knowledge acquired from such experience actually shapes online visual control remains an open question. Using a simulated driving task, we show that familiarity with a road improves steering stability specifically when visibility is reduced, and that this benefit depends on learned associations between landmarks and upcoming road geometry rather than a global cognitive map. The results indicate that spatial knowledge plays a key role in moment-to-moment control, letting drivers anticipate road segments they cannot yet see. Unfamiliar roads and impaired spatial learning may compound the risks of poor visibility, suggesting a role for driver-assistance systems that leverage landmarks.

## Introduction

Humans and other animals routinely interact with and move about through complex and dynamic environments. They reach for, grasp, and manipulate objects, locomote through crowded spaces, over rough terrain, and along winding paths, and navigate to more distant locations beyond the field of view. Such actions sometimes take place in environments that the actor has never before encountered but more often occur in environments that are familiar, in which case aspects of the spatial layout may be known. It is well established that such knowledge plays an important role in tasks that involve spatial cognition, such as finding a lost item at home, locating a car in a familiar parking lot, or finding a detour to the grocery store in a familiar town when the usual route is closed off (1-4). Whether knowledge of spatial layout also contributes to the online control of action with respect to visible objects (e.g., during visually guided steering) is an open question.

The present study was designed to better understand the possible contribution of spatial knowledge in the control of locomotion, using a simulated driving task as the context within which to investigate this issue. Specifically, the primary aim was to determine whether familiarity with the geometry of the road based on previous experience contributes to the control of steering and speed while driving along a winding road. Although this topic has not been directly addressed in previous research on driving, there are some findings that are consistent with a role for spatial knowledge. For example, Kunishige et al. (5) reported that older adults performed worse on both a spatial cognition task and a lane-changing task compared to younger adults. Within both groups, poorer spatial cognition was associated with less smooth driving. Studies of professional race car drivers provide additional evidence that spatial knowledge plays a role. Before races, drivers study the circuit to learn braking points and curve apexes (6), knowledge that may also guide their eye and head movements during the race (7).

On the other hand, people routinely drive on unfamiliar roads without incident, suggesting that knowledge of road geometry is not essential for everyday driving. In these situations, steering can be guided by currently available visual information, including the tangent point of the road (8, 9), lane edges (10), and optic flow (11-13). Consistent with this view, most computational models account for human steering using only online visual information, with no role for spatial knowledge (14-17). Moreover, the distortions and inaccuracies characteristic of spatial knowledge (3, 18-21) raise doubts about whether such knowledge is sufficiently reliable to support steering in challenging conditions such as dense fog.

Taken together, these findings paint an unclear picture about the role of spatial knowledge in steering control, motivating further research. Resolving this issue has theoretical significance insofar as locomotor control is generally understood in terms of a direct coupling of currently available information to action with no explicit role for spatial knowledge (22, 23). Finding that spatial knowledge contributes little to the control of locomotion would validate such accounts. Conversely, if spatial knowledge proves important for steering, such accounts would need further development. The issue also has practical implications, such as clarifying whether deficits in spatial cognition could impair driving (5, 24).

While the primary aim of this study was to explore the role of spatial knowledge of road geometry in the control of steering, there were also secondary and tertiary aims. The second aim was to determine whether the contribution of spatial knowledge depends on the quality of currently available visual information that is used by drivers to control steering. This aim was motivated by the possibility that knowledge of road geometry may only contribute when visibility is reduced (e.g., when driving at night or in dense fog). Lastly, spatial knowledge can take many forms (e.g., landmark-based, route knowledge, survey knowledge, cognitive maps cognitive graphs) (18, 25, 26). As such, the third aim was to better understand the nature of the spatial knowledge that drivers acquire as they repeatedly drive along the same road and use to control steering. Specifically, we aimed to determine whether such knowledge is better understood in terms of associations between landmarks and individual road segments or a global, cohesive map.

### Approach

To address these questions, we conducted two experiments in which subjects performed a simulated driving task in a desktop virtual environment, using a steering wheel and foot pedals to drive along a winding road (see Figure 1A and Movie S1). Subjects completed 10 trials, each of which involved driving along a ∼3 km winding road and took about 2-3 minutes to complete. In Experiment 1, one group (Constant Track group) followed the same track (Track A; see Figure 1B) on all ten trials. The other group (Variable Track group) followed a different track on each trial. Importantly, the track on the tenth and final trial was Track A for both groups. The visibility of the road was also manipulated within-subjects by adding dense fog to some of the road segments (see Figure 1C) while visibility was unobstructed for other segments. To address the main research question, we compared steering behavior across groups and across visibility conditions on Trial 10.

**Figure 1.**
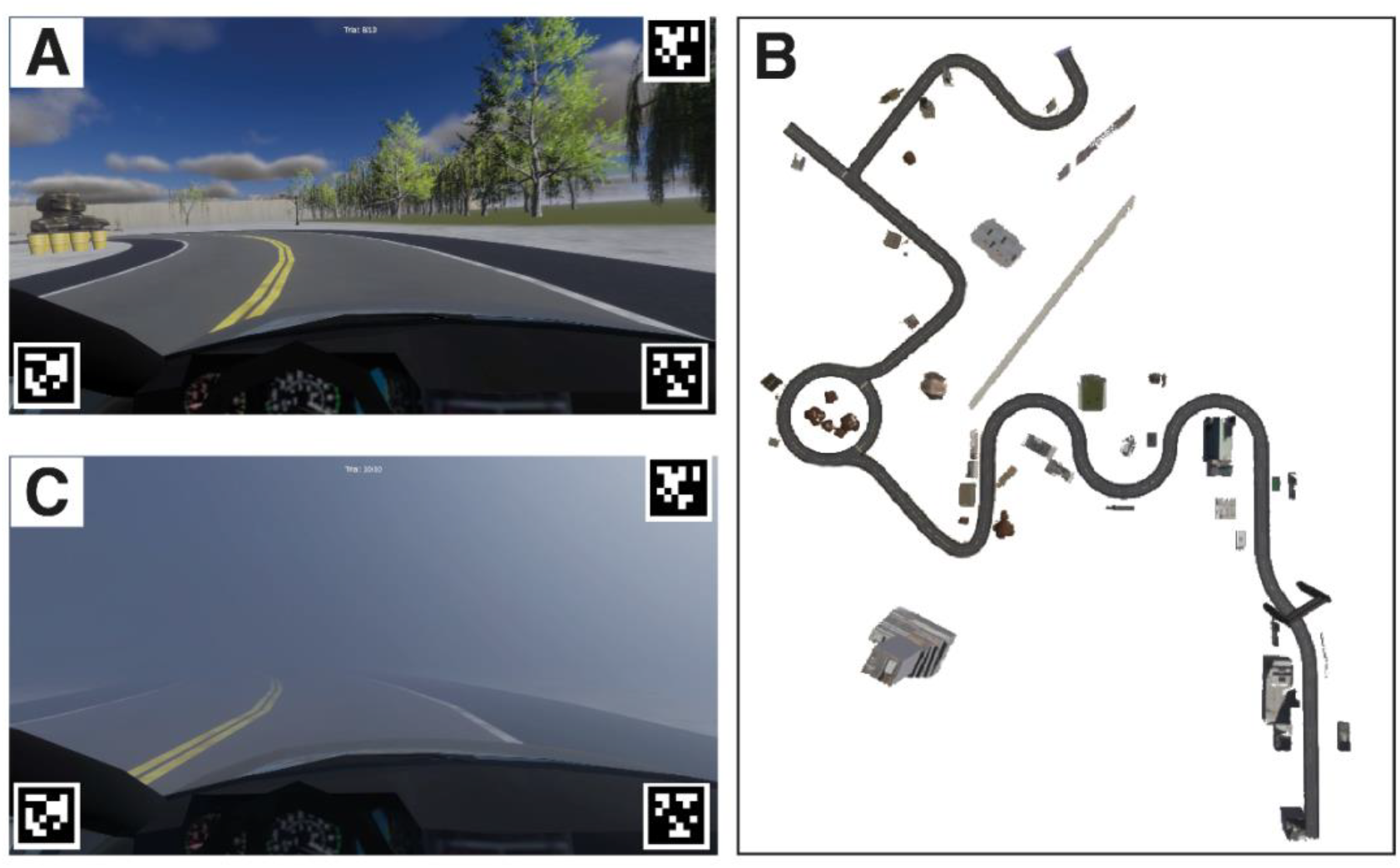
Screenshot of virtual environment (A) and top-down view (B) of Track A in Experiment 1. The straight segment in the bottom-right corner was the starting position. The elements beside the road are landmarks (e.g., buildings, rock formations). (C) Screenshot of segment of virtual environment that was surrounded by fog.

This comparison between groups assumes that when people repeatedly drive along the same road, they spontaneously learn something about the geometry of that road, even if the task does not involve any navigational decisions. In other words, it assumes that subjects in the Constant Track group learned more about the geometry of Track A than subjects in the Variable Track group. To verify this assumption, the experiments also included two post-tests that required subjects to reproduce the track from Trial 10 – one that measured subjects’ knowledge of track geometry in an egocentric reference frame and the other that measured knowledge in an allocentric reference frame. We then compared performance on the post-tests across the two groups to determine whether repeatedly driving along the same road allowed subjects to better learn the road geometry. In addition, we conducted a second experiment to understand the nature of the spatial knowledge that is used to guide high-speed steering.

## Results

### Experiment 1: Analysis of Post-test performance

The first set of analyses focuses on Post-tests 1 and 2. We begin with the post-tests rather than steering behavior to establish that subjects in the Constant Track group, who repeatedly drove along the same road (Track A), better learned the road geometry than subjects in the Variable Track group, who experienced Track A just once (on Trial 10). If driving along the same road for ten trials was not sufficient to allow subjects in the Constant Track group to learn something about the road geometry, the question about whether the two groups exhibit differences in steering performance or behavior would be moot.

Post-test 1 required subjects to reproduce the curved segments of Track A from memory using the steering wheel while viewing the virtual environment from a first-person perspective, as in the main experiment. The straight segments of the road, which alternated with the curved segments, and the landmarks were visible, but the curved segments were removed (see Figure 2A). Post-test 1 started with the vehicle at the beginning of the first straight segment. Subjects were instructed to drive along that segment until the road ended, at which point they should attempt to drive the car along a path that followed the road that they drove along in the previous trial (see Movie S2). After they drove a distance equal to the length of the curved segment, they were teleported to the beginning of the second straight segment. The task was repeated until the subject completed all eight curved segments.

**Figure 2.**
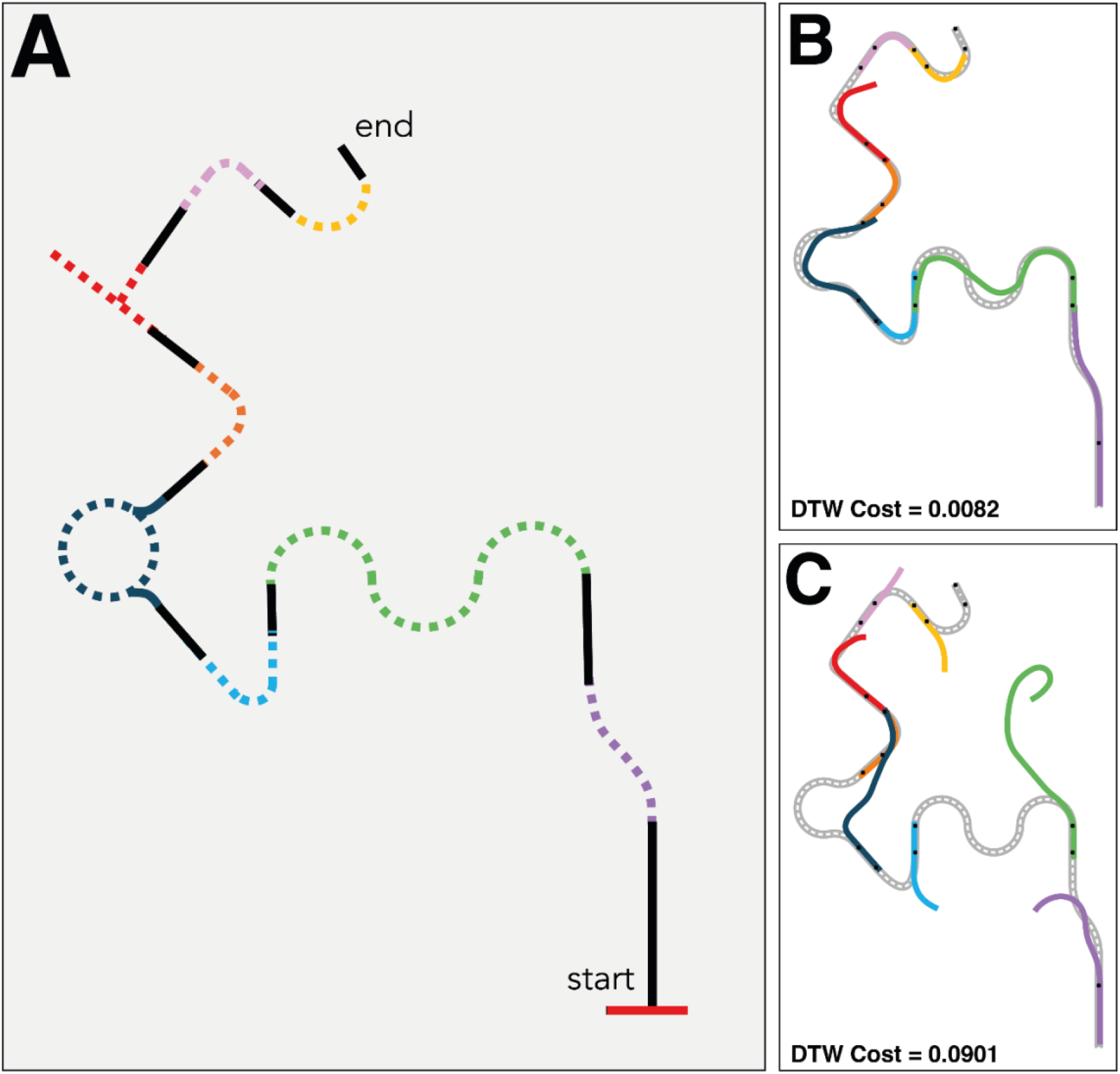
(A) Top-down view of Track A comprising straight segments (black lines) interleaved with curved segments (colored dashed lines). In Post-test 1, the straight segments and landmarks were visible, but the curved segments were not. (B and C) Examples of trajectories produced by subjects in Post-test 1 of Experiment 1, illustrating good performance (low DTW cost) and poor performance (high DTW cost).

The accuracy with which subjects reproduced the curvature of the invisible track segments was quantified by using dynamic time warping (DTW) (27, 28) to measure the similarity between the trajectory produced during Post-test 1 and the trajectory produced on Trial 10 (i.e., when the entire road was visible). DTW is an algorithm for measuring the similarity of two trajectories by finding the optimal alignment of frames that minimizes the cumulative distance between corresponding points (see Data Analysis section of Methods). Unlike other approaches, such as calculating the Euclidean distance between positions along the two trajectories at the same point in time, DTW allows for the possibility that the two trajectories could have been produced while moving at different speeds. Smaller DTW scores indicate greater similarity (see Figures 2B and 2C for examples of Post-test 1 trajectories with low and high DTW scores, respectively). Because DTW scores are bounded by zero and tend to be positively skewed, we log-transformed the raw scores prior to running an independent-samples t-test. Log-DTW scores for subjects in the Constant Track group (*M* = -4.24, 95% *CI* = [-4.60, -3.87]) were significantly less than scores for subjects in the Variable Track group (*M* = -3.60, 95% *CI* = [-4.09, -3.11]), *t*(25.95) = 2.23, *p* < .05, *g* = 0.79, 95% *CI* = [0.05, 1.51]. This indicates that the trajectories generated by subjects in the Constant Track group more closely followed the road compared to the trajectories generated by those in the Variable Track group.

Post-test 2 required subjects to hand draw an aerial map of Track A using a stylus and tablet, providing a measure of spatial knowledge in an allocentric reference frame (see Figure S1 for an example and Figure S2 in SI appendix for the entire set of drawings). Drawings were assigned accuracy scores by three independent raters, who were presented with slides, each of which showed a side-by-side view of a subject’s drawings and a top-down view of Track A. Raters were instructed to base their scores on four criteria: number of segments, geometry, order, and orientation (see Data Analysis section of Methods and SI Appendix, Section 1.B for details). An independent-samples t-test revealed that the Constant Track group (*M* = 11.56, 95% *CI* = [8.89, 14.23]) scored significantly higher than the Variable Track group (*M* = 5.58, 95% *CI* = [3.6, 7.57]), *t*(25.31) = 3.87, *p* < .001, *g* = 1.38, indicating that subjects in the Constant Track produced drawings that were judged to be more accurate depictions of the track.

Taken together, the two post-tests reveal that subjects in the Constant Track group exhibited greater recall of the track geometry. Note that we did not tell subjects that there would be a post-test or instruct them to remember anything about the shape of the road while they were performing the driving task. Furthermore, the task did not require subjects to make any navigational or route decisions – only to stay in the lane. As such, the findings of the post-test indicate that repeatedly driving along the same road was sufficient for subjects to spontaneously learn about the geometry of the road.

### Experiment 1: Analysis of steering performance

The next set of analyses focuses on whether the knowledge of road geometry that was acquired by subjects in the Constant Track group played a role in the control of steering. Specifically, we tested for differences between the Constant Track group and Variable Track group across various measures of steering performance on Trial 10. Because the contribution of spatial knowledge may depend on the availability of visual information, we also included road visibility as an independent variable, comparing performance on clear segments versus foggy segments. The effects of track constancy and visibility were assessed on four dependent measures of steering performance: mean speed, variability in speed, mean steering wheel angular acceleration, and variability in lane position. For each measure, we ran a two-way (track constancy x visibility), mixed-design ANCOVA with mean steering wheel acceleration on Trial 1 as a covariate. The covariate was included to control for baseline differences in driving skill between subjects.

Not surprisingly, subjects drove significantly faster in High Visibility segments compared to Low Visibility segments [*F* (1,27) = 411.5, *p* < 0.001, η^2^_p_ = 0.938]. However, neither the main effect of track constancy [*F* (1, 27) = 0.49, *p* = 0.49, η^2^_p_ = 0.018] nor the track constancy x visibility interaction [*F* (1, 27) = 0.031, *p* = 0.861, η^2^_p_ = 0.001] was significant (see Figure 3A). A similar pattern of results was observed in the analysis of speed variability. Subjects maintained a more consistent speed (i.e., exhibited lower speed variability) in the High Visibility segments [*F*(1, 27) = 56.938, *p* < 0.001, η^2^_p_ = 0.678] but the main effect of track constancy [*F*(1, 27) = 2.478, *p* = 0.127, η^2^_p_ = 0.084] and the track constancy x visibility interaction [*F*(1, 27) = 0.348, *p* = 0.56, η^2^_p_ = 0.013] were not significant (see Figure 3B).

**Figure 3.**
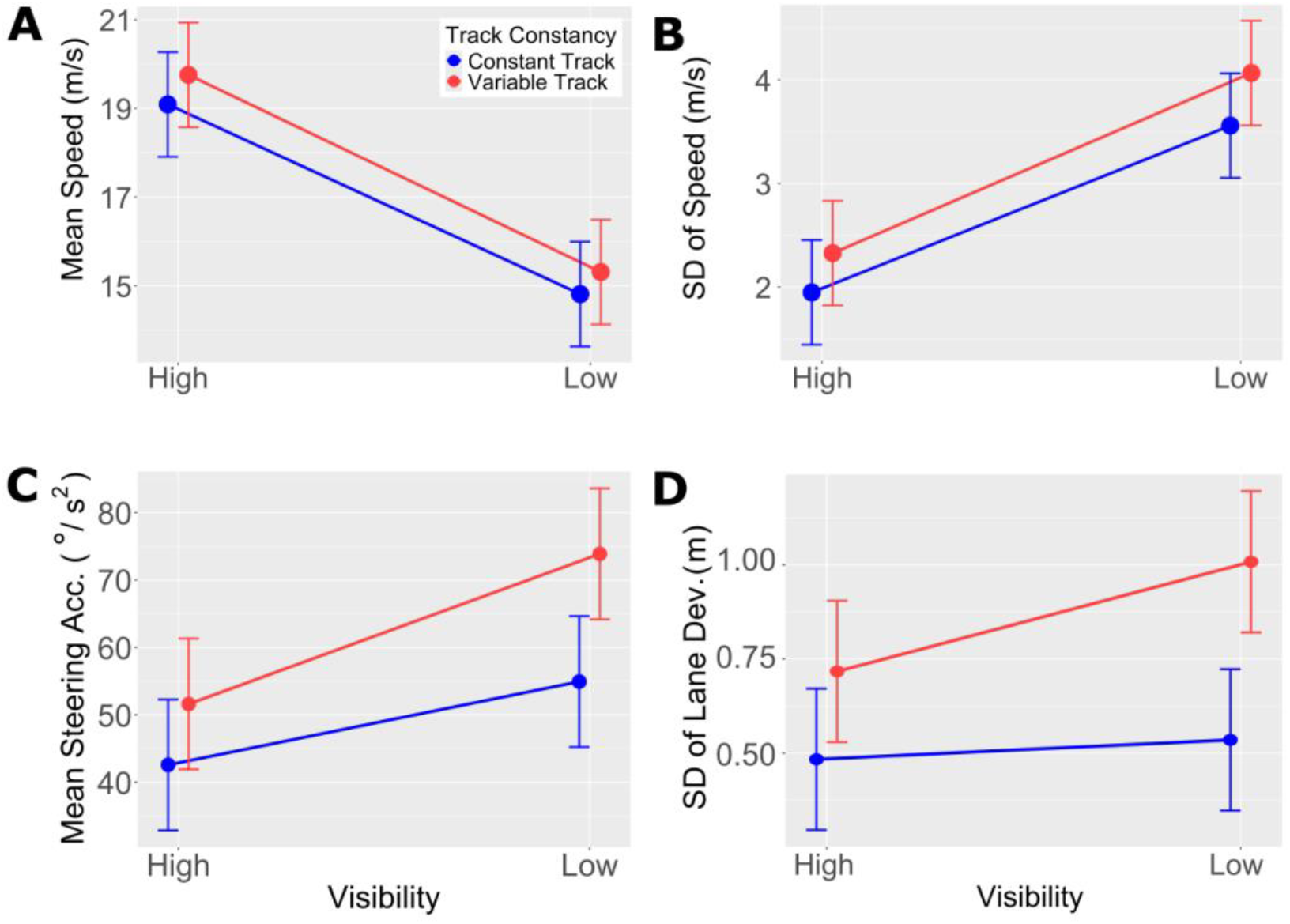
Mean speed (A) standard deviation of speed (B) mean absolute steering wheel acceleration (C) and standard deviation of lane position (D) as a function of road visibility on Trial 10 of Experiment 1 for subjects in the Constant Track (blue) and Variable Track (red) groups. Error bars indicate 95% confidence intervals.

Interestingly, the analysis of mean steering wheel angular acceleration revealed a significant track constancy x visibility interaction [*F*(1,27) = 4.693, *p* < 0.05, η^2^_p_ = 0.148; see Figure 3C]. Post-hoc pairwise comparison with Bonferroni adjustment showed that angular acceleration was greater for the Variable Track group compared to the Constant Track group in Low Visibility segments (*p* < 0.01). However, this difference did not reach statistical significance in High Visibility segments (*p* = 0.198).

A similar pattern of results was observed in the analysis of lane position variability [*F*(1,27) = 4.472, *p* < 0.05, η^2^_p_ = 0.142; see Figure 3D]. Variability was significantly greater in the Variable Track group compared to the Constant Track group in Low Visibility segments (*p* < 0.001) but not in High Visibility segments (*p* = 0.09). Taken together, these results suggest that when subjects were familiar with the track, they were better able to control steering but only under low visibility conditions.

If in fact, knowledge of road geometry contributed to the control of steering, then subjects who scored higher on the post-tests should also exhibit better steering performance. Indeed, performance on Post-test 1, in which subjects reproduced the curved segments of the track from a first-person perspective, was significantly correlated with both steering acceleration (*r*_*s*_ = .65, *p* < .001, *n* = 30; see Figure S3A in SI Appendix) and lane position variability (*r*_*s*_ = .707, *p* < .0001, *n* = 30; see Figure S3B in SI Appendix) in road segments that were surrounded in fog. However, we did not find significant correlations involving Post-test 2, in which subjects produced hand-drawings of the track from an aerial perspective (*rs* = -.11, *p* = 0.58, *n* = 29 for steering acceleration and *rs* = -.087, *p* = 0.65, *n* = 29 for lane position variability; see Figures S3C and S3D, respectively. in SI Appendix). This suggests that while repeatedly driving along the same road was sufficient to learn about the road geometry in both egocentric and allocentric reference frames, the control of steering may have relied only on spatial knowledge in an egocentric rather than allocentric reference frame. Experiment 2 was designed to further explore this hypothesis.

### Experiment 2

Having established that spatial knowledge plays a role in the control of high-speed steering (albeit only when visibility is reduced), we turn to the question of the nature of such knowledge. We considered two hypotheses that differ in terms of whether drivers rely on local associations between landmarks and road segments or global, route-based or map-like knowledge. The distinction shares elements of the contrast between landmark, route, and survey knowledge that was drawn by Siegel and White (26) in their seminal work on the acquisition of spatial knowledge. In the present study, however, we focus on the distinction between the one form of knowledge that is local (landmark-based knowledge) and the two forms of knowledge that are global (route-based and survey knowledge).

The local associations hypothesis is that when people repeatedly drive along the same road, they learn local associations between salient landmarks and individual road segments (e.g., there is a sharp turn to the right immediately after the post office). Such knowledge is fragmented in the sense that it does not encode the sequential order, relative distances, or relative directions of road segments. The global knowledge hypothesis is that drivers integrate knowledge about the environment into a cohesive, global map that preserves the sequential order of road segments (e.g., after turning sharply to the right, go straight for a short period before entering a long, gradual turn to the left) or perhaps even the geometric structure, relative distances, and relative directions of road segments.

Previous research indicates that landmarks are salient visual cues used for guiding navigation decisions in the context of driving, in tasks such as route selection and planning (29-32). However, it is not clear whether landmarks may also be used in the online control of steering and speed to maintain lane position. Furthermore, there is also a question of whether landmarks are solely used to guide the visual control of steering, or if landmarks instead aid the development of global cognitive maps of the track that guide high-speed steering. Lastly, the analysis from Experiment 1 indicating a correlation between steering performance and Post-test 1 but not Post-test 2 (see Figure S3 in SI Appendix) is consistent with the local associations hypothesis, as Post-test 1 required subjects to employ an egocentric reference frame whereas Post-test 2 required subjects to employ an allocentric reference frame. However, this finding is far from conclusive, motivating the need for a more definitive test of whether the knowledge that drivers use to improve the control of steering is best characterized as local or global.

We manipulated the configuration of track segments and landmarks as a between-subjects variable with three groups (see Figure 4). In the Control group, subjects drove along the same track with the same landmark arrangement for all ten trials, similar to the Constant Track group in Experiment 1. In this condition, both local landmark associations and the global structure of the track were preserved. In the Scrambled Landmarks group, the track was the same on all ten trials, but the landmarks appeared in different locations. For example, the building that was in the center of the circular segment on one trial was replaced with a boulder on another trial, and the building appeared next to a different track segment (see middle column of Figure 4). This condition also preserved global structure, but local landmark-road segment associations were manipulated. In the Scrambled Segments group, subjects drove along a different track in each of the ten trials, with tracks comprising the same set of track segments in different orders. Importantly, individual landmarks remained in the same positions relative to individual segments (see right column of Figure 4). Hence, the global structure was manipulated while local associations were preserved. As in Experiment 1, the track that subjects followed on the 10th and final trial was the same across all three groups.

**Figure 4.**
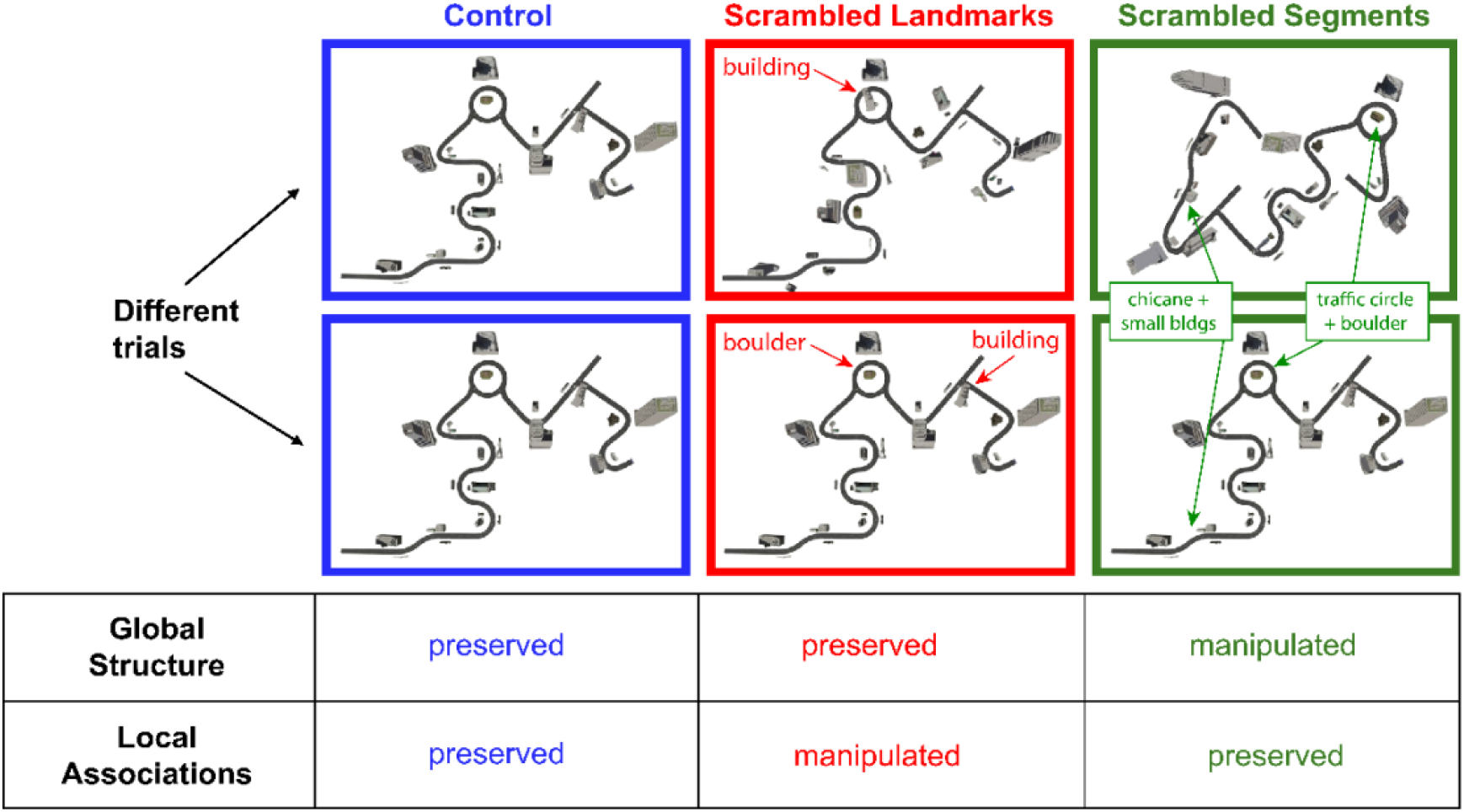
Three conditions in Experiment 2. In the Control condition, the track and arrangement of landmarks were the same for all ten trials, preserving both the global structure and local associations. In the Scrambled Landmarks condition, the track was the same on each trial, but the landmarks appeared in different positions. This condition preserved the global structure but manipulated the local associations. In the Scrambled Segments condition, the road segments appeared in different positions on each trial, but the landmarks remained in the same positions relative to the road segments. As such, global structure was manipulated but the local associations were preserved.

If drivers rely on local associations between landmarks and track segments to control steering, performance in the Scrambled Landmarks group should be worse compared to performance in the Scramble Segment and Control groups. On the other hand, if drivers rely on global knowledge of the track to guide steering, performance should be worse in the Scrambled Segments group. Based on the results of Experiment 1, we expect any differences across groups to be most pronounced in the low-visibility segments.

### Experiment 2: Analysis of Post-test performance

As in Experiment 1, we start with the analyses of Post-tests 1 and 2 to determine whether subjects’ ability to recollect the spatial layout of the track on Trial 10 was affected by the manipulation of track configuration. Mean log-DTW scores were higher for subjects in the Scrambled Landmarks (*M* = -3.73, 95% *CI* = [-4.01, -3.45]) and Scrambled Segments (*M* = -3.82, 95% *CI* = [-4.02, -3.62]) groups compared to those in the Control group (*M* = -4.134, 95% *CI* = [-4.44, -3.83). Although the effect did not reach statistical significance, *F* (2,57) = 2.83, *p* = 0.07, η^2^_p_ = 0.09, closer inspection revealed that mean log-DTW cost was highest in the Scrambled Landmarks condition for six of the eight track segments.

For Post-test 2, a one-way independent-samples ANOVA revealed a significant effect of track configuration on the accuracy scores assigned to drawings, *F*(2,57) = 5.376, *p* < 0.01, η^2^_p_ = 0.159 (see Figures S4 and S5 in the SI Appendix). Post-hoc pairwise comparison with Bonferroni adjustment showed that the Scrambled Segments group had significantly lower accuracy scores (*M* = 8.27, 95% *CI* = [7.01, 9.53]) than the Control group (*M* = 11.7, 95% *CI* = [10.2, 13.2]), *p* < 0.05). Accuracy scores in the Scrambled Landmarks group (*M* =10.4, 95% *CI* = [8.78, 12.0]) were not significantly different from those in the Control group.

### Experiment 2: Analysis of steering performance

To better understand the nature of spatial knowledge that is used to guide high-speed steering, we tested differences in speed, steering, and lane position control across the three groups on Trial 10. As in Experiment 1, subjects drove faster [*F* (1, 56) = 370.25, *p* < 0.001, η^2^_p_ = 0.869] and exhibited lower speed variability [*F*(1, 56) = 266.73, *p* < 0.001, η^2^_p_ = 0.826] in segments with high visibility. However, neither the effect of track configuration [*F* (2, 56) = 0.620, *p* = 0.524, η^2^_p_ = 0.022 for mean speed; *F*(2, 56) = 0.004, *p* = 0.996, η^2^_p_ = 0.0002 for speed variability] nor the track configuration x visibility interaction [*F*(2, 56) = 0.157, *p* = 0.855, η^2^_p_ = 0.006 for mean speed; *F*(2, 56) = 0.667, *p* = 0.517, η^2^_p_ = 0.023 for speed variability] were significant for either speed-related dependent measure (see Figure 5A and 5B].

**Figure 5.**
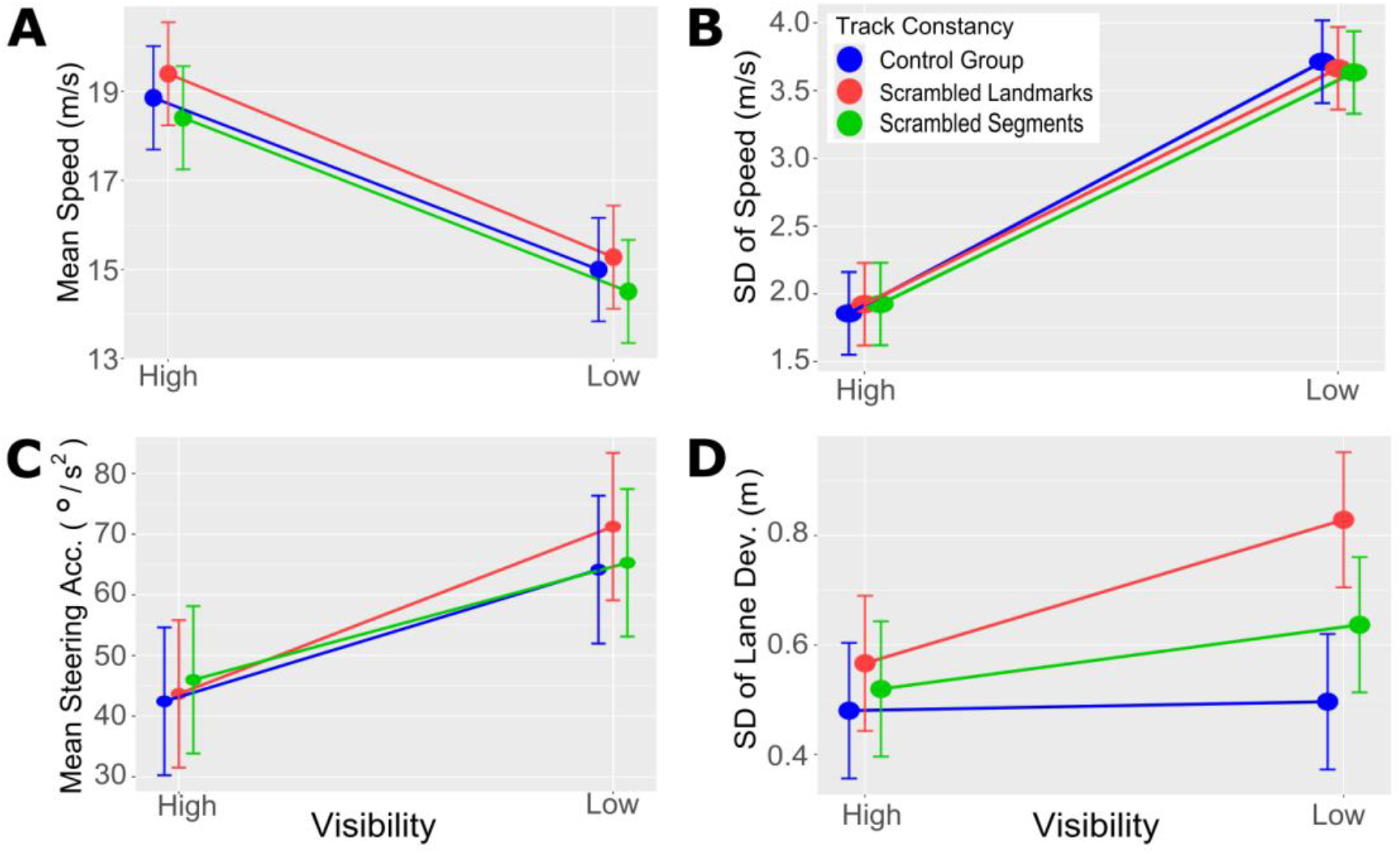
Mean speed (A) standard deviation of speed (B), mean absolute steering acceleration (C) and standard deviation of lane position (D) on Trial 10 plotted as a function of visibility (High vs. Low) on Trial 10 OF Experiment 2 for subjects in the Control, Scrambled Landmarks, and Scrambled Segments conditions. Error bars indicate 95% confidence intervals.

The analysis of steering wheel angular acceleration also revealed a significant main effect of visibility [*F*(1,56) = 46.476, *p* < 0.01, η^2^_p_ = 0.454], with higher mean accelerations in Low Visibility segments (see Figure 5C). However, neither the main effect of configuration [*F*(2,56) = 0.154, *p* = 0.858, η^2^_p_ = 0.005] nor the configuration x visibility manipulation [*F*(2,56) = 0.524, *p* = 0.595, η^2^_p_ = 0.018] were significant, suggesting that steering stability was consistent across the three groups.

The analysis of lane position variability revealed significant main effects of both visibility [F(1,56) =13.302, p < 0.01, η^2^_p_ = 0.192] and track configuration [*F*(2,56) = 3.769, *p* < 0.05, η^2^_p_ = 0.119], as well as a significant track configuration x visibility interaction [*F*(2,56) = 3.296, *p* < 0.05, η^2^_p_ = 0.105; see Figure 5D]. Post hoc pairwise comparisons with Bonferroni adjustment revealed that, under low visibility, lane position variability was significantly greater in the Scrambled Landmarks group compared to the Control group, *p* < .001. However, the Scrambled Segments group was not significantly different (*p* = .115) than the Control group. Under high visibility conditions, no significant differences were observed (*p* > .05) across any pairs of groups. Taken together, the results are most consistent with the hypothesis that drivers rely on local associations between landmarks and track segments. In addition, the findings reinforce those from Experiment 1 suggesting that knowledge of road geometry only plays a meaningful role under reduced visibility conditions. When visibility is good, such knowledge plays a negligible role.

## Discussion

### Summary of results

The main findings of the present study can be distilled into three points. First, participants who repeatedly drove along a road showed better knowledge of its layout than those who drove it only once, as measured by both a first-person steering task and a bird’s-eye sketch of the road. This was true even though participants were not forewarned that they would be tested on their knowledge of the road and even though the driving task itself required no navigational decisions, such as whether to turn left or right at an intersection. This suggests that repeatedly driving along the same road is sufficient to learn about the geometry of the road in both egocentric and allocentric reference frames and that such learning occurs incidentally as a byproduct of following the route.

Second, participants who repeatedly drove along the same track exhibited better performance on measures of steering stability (e.g., mean absolute steering acceleration and standard deviation of lane position). However, the benefits of familiarity were modest and only observed under low-visibility conditions. This finding attests to the primacy of currently available visual information. When visual information that is sufficient to perform the task is available, prior knowledge contributes very little to the control of action. On the other hand, such knowledge can help sustain steering stability when it would otherwise deteriorate, allowing for the control of action to remain robust across a range of visibility conditions.

Third, steering performance was surprisingly robust to the scrambled segments manipulation, which affected the ordering of road segments across trials but maintained the relative positions of individual landmarks to road segments. Conversely, when visibility was reduced, lane position variability increased under the scrambled landmarks manipulation, which left the ordering of road segments intact but affected the relative positions of landmarks to road segments. This pattern of results suggests that the knowledge of road layout upon which drivers rely to maintain steering stability under reduced visibility conditions is best characterized in terms of local associations between landmarks and road segments. While drivers may also acquire knowledge of the road’s global structure, such knowledge does not appear to be used to control steering. Consistent with this interpretation, there was a positive correlation between steering stability and performance on Post-test 1, which measures knowledge of road geometry in egocentric coordinates. However, no such correlation was found for Post-test 2, which reflects knowledge in allocentric coordinates.

### Theoretical implications

In the section, we consider how these findings can be reconciled with the conventional view that currently available information is sufficient to control steering. First, it is worth emphasizing that road familiarity only influenced steering performance when visual information was degraded. In high visibility conditions, participants who were more familiar with the road performed no better than those for whom the road geometry was unfamiliar. This finding reaffirms rather than challenges one of the central claims of the information-based account (22, 23) – that under normal viewing conditions, currently available information is sufficient to perform the task without the need for prior knowledge.

The finding that road familiarity selectively affected steering stability provides a potentially important clue as to how spatial knowledge may contribute to steering control. A well-established distinction in the steering literature is between anticipatory guidance control, which uses information from more distant regions of the road to generate smooth, prospective adjustments in anticipation of an upcoming curve, and compensatory stabilization control, which uses information from nearby regions to correct lane position deviations (17, 33, 34). Steering stability in particular is thought to reflect the quality of anticipatory control, as steering tends to be smoother when drivers can preview upcoming road curvature from a distance compared to when they must react to each new segment of a road as it arrives (8, 35). Under reduced visibility conditions, information from more distant regions of the road is unavailable and anticipatory control is compromised. The present findings suggest that under these conditions, landmarks associated with specific road segments can serve as information about the upcoming road curvature that is needed for anticipatory steering control. Just as information from more distant regions of the road specifies upcoming curvature under good viewing conditions, a familiar landmark can inform the driver about the steering adjustments that will soon be required.

This account could explain why the knowledge of road geometry that participants in the present study used to control steering was landmark-based rather than map-like. When a distinctive landmark remains in a fixed position relative to an individual road segment, that landmark serves as a reliable predictor of the upcoming steering demands; that is, the landmark carries information for the control of steering much like road edges and optic flow. The driver simply needs to become attuned to this regularity as they repeatedly encounter that section of the road. This kind of learning, in which an arbitrary perceptual cue is recruited to support the control of action, has empirical precedent. Li, Saunders, and Li (36) showed that participants performing a manual tracking task learned to use a novel color cue that correlated with target motion to improve their performance, suggesting that the visual-motor system can become attuned to any reliable cue, however arbitrary, provided it carries information relevant to the control demands of the task. From this perspective, landmarks provide information about the upcoming action (i.e., steering adjustment) rather than the geometry of the road per se. This interpretation is consistent with the finding that steering performance in Experiment 1 was correlated with performance on Post-test 1 but not Post-test 2.

Thus, we propose the contribution of spatial knowledge to steering control is best understood not as the retrieval of a stored map, but as the use of learned landmark-road curvature associations to maintain anticipatory control when information from more distant road regions is unavailable. More broadly, this interpretation is compatible with the emerging view (3, 21, 37, 38) that spatial knowledge that is fragmented and landmark-anchored is sufficient to account for many aspects of navigation that were previously assumed to require a cognitive map (39, 40).

### Practical significance

Although the broader significance of the present study is primarily theoretical, the findings also carry some practical implications for understanding the factors that contribute to driving risk. The fact that road familiarity improved steering stability specifically under reduced visibility conditions suggests that familiarity and visibility interact in ways that compound risk. Specifically, reduced visibility alone degrades steering stability, but the effect is more pronounced when the environment is unfamiliar. While it is well known that driving in fog, rain, or at night increases risk, the present findings suggest that this risk may be elevated when drivers are on unfamiliar roads, where they haven’t yet learned landmark-road segment associations that could play a role when poor visibility disrupts the detection of distant visual information.

The findings also raise questions about the risks faced by drivers with diminished spatial learning or memory abilities, such as older adults or individuals in the early stages of neurodegenerative conditions (5, 24). If such drivers are less able to learn the kinds of landmark-road segment associations that support anticipatory steering under reduced visibility, they may be at elevated risk in conditions where those associations would otherwise compensate for degraded visual information. Use of advanced driver assistive systems that provide augmented reality overlays or that highlight landmarks to promote learning (41) could be especially critical in mitigating driving risk under these conditions.

## Conclusion

The present findings demonstrate that spatial knowledge acquired through repeated experience with a road can support the online control of steering, albeit only when currently available visual information is insufficient to do so. Under such conditions, the benefit of familiarity appears to derive from learned associations between local landmarks and road geometry rather than from a globally coherent representation of the route. The role of spatial knowledge is not limited to the kinds of off-line, memory-guided tasks with which it has traditionally been associated, such as navigating to an unseen destination, finding a lost object, or planning a detour, but extends to the online visual control of action in familiar environments. Future research should determine how these landmark associations are acquired, including the influence of landmark placement, gaze behavior, cognitive load, and individual differences in spatial cognition, and should extend existing steering models (e.g., 16, 17) to incorporate learning mechanisms that use landmark associations to support anticipatory steering under reduced visibility.

## Materials and Methods

### Participants

Thirty subjects [15 women, 15 men; mean age = 19.74 (18-26) years] participated in Experiment 1, and sixty subjects [20 women, 37 men; mean age = 20.4 (18-30) years] participated in Experiment 2. Data from one subject in Experiment 1 was included in the steering performance analysis but excluded from the post-test analyses because the subject took unusually long to complete trials 1–10 and had to leave before completing the post-tests. Relevant self-reported measures such as handedness, driving experience, and video game play for participants in both experiments are provided in Table S1 and Table S2 in SI appendix. Participants in Experiment 1 were recruited from psychology courses and received extra credit for participation, and participants in Experiment 2 were recruited from the university campus and received $15 for participation. None of the participants reported any motor impairments, and all reported normal or corrected-to-normal vision. The use of an eye tracker in Experiment 1 precluded subjects from wearing eyeglasses; accordingly, all subjects had either uncorrected normal vision or corrected-to-normal vision using contact lenses. The protocol was approved by the Institutional Review Board at Rensselaer Polytechnic Institute, and all subjects gave informed consent before performing the experiment.

### Hardware

The experiment was run on an Alienware Aurora R15 (Dell, Round Rock, TX) equipped with an NVIDIA GeForce RTX 4090 graphics card (NVIDIA, Santa Clara, CA) and a 3.0-GHz 13th Generation Intel Core i9-13900KF processor (Intel, Santa Clara, CA). The scene was presented to subjects on a 55-inch LG 55UF85 monitor (Seoul, South Korea) with a 1920 × 1080 resolution at 60 Hz. Steering and speed were controlled using a Logitech G920 Driving Force Racing Wheel and Pedals system. In Experiment 1, subjects’ eye movements were tracked using a Pupil Labs Core eye tracker (Berlin, Germany), although the eye tracking data were not analyzed.

### Virtual environment

Ten unique tracks were created for Experiment 1 and labeled A through J. Each track consisted of eight straight segments alternating with eight curved segments (see Figure 2 for an example). For the nine tracks used for trials 1 through 9 in the Variable Track group, the curved segments were randomly selected from a set of 11 segment shapes, including horseshoes, a zig-zag, a U-turn, and a lane convergence. For Trial 10, the track consisted of a different set of eight curved segments, including turns to the right or left of varying curvatures and arc lengths, a traffic circle, a chicane, a triple S-turn, and a T-intersection. This track was also used on every trial for the Constant Track group. In Experiment 2, all tracks across all three conditions were constructed from this same set of curved segments.

Tracks were given a light-gray asphalt texture and divided into two 5 m-wide lanes by a double-yellow centerline (see Figure 1A). The outer edge of each lane was marked by a single white line and bordered by a narrow shoulder with a darker-gray asphalt texture. During Trials 4, 6, and 7, the track included fog on a single curved segment so that participants could become familiar with driving under reduced visibility conditions. On Trial 10, three of the curved track segments were surrounded by fog (Low Visibility) while the remaining five curved segments were not (High Visibility).

Each track was embedded within a unique virtual scene comprising a combination of two regions, including human-made regions (e.g., urban, suburban) and more natural regions (e.g., beach, forest, desert). Each region was populated with objects commonly associated with that environment (e.g., urban regions contained buildings, suburban regions contained houses and fences, forest regions contained trees and rock formations). In addition to giving each region a distinct visual character, these objects provided landmarks that afforded the possibility of learning associations with the upcoming road segment.

In Experiment 2, the track that was used on all ten trials for subjects in the Control and Scrambled Landmarks groups was the same as Track A from Experiment 1. Track A was also used for the tenth and final trial for subjects in the Scrambled Segments group. The tracks used for the other nine trials in the Scrambled Segments group were created by rearranging the segments from Track A.

The surrounding environments in Experiment 2 did not contain regions as in Experiment 1 because regions would have been disrupted by the scrambling of segments and landmarks. Instead, landmarks were arranged in clusters surrounding the track and were present in all conditions but differed in their consistency across segments: clusters were fixed to specific segments for the Control and Scrambled Segments groups, whereas in the Scrambled Landmarks group the same clusters appeared with different segments across Trials 1–9.

The vehicle (a sedan) was an asset purchased from the Unity Asset Store (Edy’s Vehicle Physics, https://assetstore.unity.com/packages/tools/physics/edy-s-vehicle-physics-403) and was controlled by the steering wheel and two foot pedals (an accelerator and a brake pedal). Subjects were able to control speeds from 0 to 60 mph (∼27 m/s). Subjects viewed the environment through the windshield of the vehicle, with the camera positioned in the driver’s seat. The dashboard and the upper part of a virtual steering wheel were visible to subjects.

### Design and procedure

Experiment 1 was a 2 × 2 mixed design in which track constancy was manipulated between subjects (Variable Track vs. Constant Track) and visibility was manipulated within subjects (Low vs. High Visibility). Participants in the Variable Track group were presented with a randomized sequence of Tracks B through J followed by Track A. Subjects in the Constant Track group were presented with Track A on all ten trials. Low Visibility segments were surrounded by fog, while High Visibility segments did not have fog. Experiment 2 was a 3 × 2 mixed design, with track configuration manipulated between subjects (Control, Scrambled Landmarks, and Scrambled Segments groups) and visibility manipulated within subjects using the same Low and High Visibility manipulation as in Experiment 1.

Prior to the start of the experiment, subjects signed a written consent form and were presented with slides containing instructions for the experiment. Next, they completed two practice sessions to become familiar with the controls and the task. The first practice session required subjects to complete five laps on a simple square-shaped track with rounded corners. In the second practice session, subjects completed three laps on a 3-km track composed of segments used in the main experiment with a different arrangement. Subjects then completed ten experimental trials, each lasting approximately three minutes, in which they drove along a track while attempting to stay in the center of the right lane, adjusting speed as necessary to maintain lane position (e.g., slowing for tight turns). Following the experimental trials, subjects completed two post-tests. Post-test 1 assessed participants’ ability to reproduce segments of the track that they followed in Trial 10 from memory from a first-person perspective (i.e., an egocentric reference frame). Subjects drove along the same track that was used in Trial 10 except that the curved segments were removed. The straight segments and the landmarks remained visible. Segments were completed one at a time. After the subject drove a distance equal to the length of the curved segment, the simulation paused and the vehicle was teleported to the beginning of the next straight segment of the track. This process was repeated until subjects completed all eight segments. In Post-test 2, participants used a tablet and stylus to sketch an aerial view of the Trial 10 track, including landmarks. This task utilized sketch mapping to help participants externalize their cognitive maps in an allocentric reference frame (42).

### Data analyses

The first set of analyses focused on results from the two post-tests. For Post-test 1, the accuracy of trajectories that subjects generated was assessed using Dynamic Time Warping (DTW). Specifically, we used DTW to quantify the geometric similarity between trajectories on curved segments of Post-test 1, in which the road was not visible, and trajectories in Trial 10, when the road was visible. Additional details about the DTW analysis are provided in the SI Appendix, Section 1.A. For Post-test 2, a rating system was developed to qualitatively analyze the geometric accuracy of each drawing (see SI Appendix, Section 1.B for details). Three raters were instructed (in individual sessions) to view each drawing alongside the ground truth outline of the track and assign a score on a scale between 0 to 5 for each criterion, with 0 indicating that no segments in the drawing matched the criterion and 5 indicating that all five segments in the drawing matched the criterion. Once ratings were completed, a composite score for each drawing was computed by summing each score across the four criteria. Then, the mean was computed for the composite scores across the three raters for each drawing. The effects of track constancy on mean log-transformed DTW scores from Post-test 1 and mean ratings from Post-test 2 were assessed using Welch’s t-tests, and effect sizes were quantified using Hedges’ g.

The second set of analyses focused on the effects of track constancy (Experiment 1) and track configuration (Experiment 2) as well as visibility on four measures of steering performance on Trial 10: mean speed, standard deviation of speed, mean absolute angular acceleration of the steering wheel, and standard deviation of lane position (see SI Appendix, Section 1.C for details about the calculation of these measures). The effects of group and visibility on each of these four measures was assessed using a mixed-design ANCOVA with mean steering wheel acceleration on Trial 1 as a covariate. The covariate was included to control for baseline differences in performance between subjects.

We conducted an additional analysis to examine the relationship between post-test performance and steering performance on Trial 10. Specifically, the mean log-DTW cost from Post-test 1 across both the Constant and Variable Track groups and the mean drawing score from Post-test 2 were compared with two steering performance metrics: lane deviation variability and mean steering acceleration under the Low Visibility condition. Steering acceleration and lane deviation variability were the only selected performance measures for this analysis because these were the only measures influenced by spatial knowledge of the track under low visibility conditions.

## Acknowledgments and funding sources

The authors thank Kiran Koch-Matthews and Xavier Marshall for developing the driving simulator and Louis Singh for assistance with the DTW analysis. This research was supported by a grant from the National Science Foundation (NSF BCS 2218220) to BRF and an RPI HASS Student Creativity and Research Award to GCR. GCR was partially supported by a National Defense Science and Engineering Graduate Fellowship.

## References

1. Epstein RA, Patai EZ, Julian JB, Spiers HJ, The cognitive map in humans: spatial navigation and beyond. Nature Neuroscience 20(11):1504–1513 (2017).

2. Golledge RG Wayfinding behavior: Cognitive mapping and other spatial processes (JHU press, Baltimore, 1999).

3. Peer M, Brunec IK, Newcombe NS, Epstein RA, Structuring knowledge with cognitive maps and cognitive graphs. Trends in Cognitive Sciences 25(1):37–54 (2021).

4. Wolbers T, Hegarty M, What determines our navigational abilities? Trends in Cognitive Sciences 14(3):138–146 (2010).

5. Kunishige M et al., Spatial navigation ability is associated with the assessment of smoothness of driving during changing lanes in older drivers. Journal of Physiological Anthropology 39(1):25 (2020).

6. Lappi O, Dove A The Science of the Racer’s Brain (2022).

7. Land MF, Tatler BW, Steering with the head: The visual strategy of a racing driver. Current Biology (2001).

8. Land MF, Lee DN, Where we look when we steer. Nature 369(6483):742–744 (1994).

9. Lappi O, Future path and tangent point models in the visual control of locomotion in curve driving. Journal of Vision 14(12):21–21 (2014).

10. Kountouriotis GK, Floyd RC, Gardner PH, Merat N, Wilkie RM, The role of gaze and road edge information during high-speed locomotion. Journal of Experimental Psychology: Human Perception and Performance 38(3):687 (2012).

11. Kountouriotis GK et al., Optic flow asymmetries bias high-speed steering along roads. Journal of Vision 13(10):23–23 (2013).

12. Kountouriotis GK, Mole CD, Merat N, Wilkie RM, The need for speed: global optic flow speed influences steering. Royal Society Open Science 3(5):160096 (2016).

13. Mole CD, Kountouriotis G, Billington J, Wilkie RM, Optic flow speed modulates guidance level control: New insights into two-level steering. Journal of Experimental Psychology: Human Perception and Performance 42(11):1818–1838 (2016).

14. Kolekar S, De Winter J, Abbink D, Human-like driving behaviour emerges from a risk-based driver model. Nature Communications 11(1):4850 (2020).

15. Lappi O, Mole C, Visuomotor control, eye movements, and steering: A unified approach for incorporating feedback, feedforward, and internal models. Psychological Bulletin 144(10):981–1001 (2018).

16. Markkula G, Boer E, Romano R, Merat N, Sustained sensorimotor control as intermittent decisions about prediction errors: computational framework and application to ground vehicle steering. Biological Cybernetics 112(3):181–207 (2018).

17. Salvucci DD, Gray R, A two-point visual control model of steering. Perception 33(10):1233– 1248 (2004).

18. Ishikawa T, Montello DR, Spatial knowledge acquisition from direct experience in the environment: Individual differences in the development of metric knowledge and the integration of separately learned places. Cognitive Psychology 52(2):93–129 (2006).

19. Tversky B, Distortions in memory for maps. Cognitive Psychology 13(3):407–433 (1981).

20. Tversky B, Distortions in cognitive maps. Geoforum 23 (2):131–138 (1992).

21. Warren WH, Non-Euclidean navigation. Journal of Experimental Biology 222(Pt Suppl 1)(2019).

22. Warren WH, Visually controlled locomotion: 40 years later. Ecological Psychology 10(3-4):177–219 (1998).

23. Zhao H, Warren WH, On-line and model-based approaches to the visual control of action. Vision Research 110(PB):190–202 (2015).

24. Kunishige M, Fukuda H, Iida… T, Spatial navigation ability and gaze switching in older drivers: A driving simulator study. Hong Kong Journal of Occupational Therapy 32:22–31 (2019).

25. Montello DR, in Spatial and Temporal Reasoning in Geographic Information Systems, eds Egenhofer MJ, Golledge RG (Oxford University Press, New York), pp 143–154 (1998).

26. Siegel AW, White SH, The development of spatial representations of large-scale environments. Advances in Child Development and Behavior 10:9–55 (1975).

27. Salvador S, Chan P, Toward accurate dynamic time warping in linear time and space. Intelligent data analysis 11(5):561–580 (2007).

28. Taylor J, Zhou X, Rouphail NM, Porter RJ, Method for investigating intradriver heterogeneity using vehicle trajectory data: A dynamic time warping approach. Transportation Research Part B: Methodological 73:59–80 (2015).

29. Edwards SJ, Emmerson C, Namdeo A, Blythe PT, Guo W, Optimising landmark-based route guidance for older drivers. Transportation Research Part F: Traffic Psychology and Behaviour 43:225–237 (2016).

30. Epstein RA, Vass LK, Neural systems for landmark-based wayfinding in humans. Philosophical Transactions of the Royal Society B: Biological Sciences 369(1635):20120533 (2014).

31. May AJ, Ross T, Presence and quality of navigational landmarks: effect on driver performance and implications for design. Human Factors 48(2):346–361 (2006).

32. Yesiltepe D, Conroy Dalton R, Ozbil Torun A, Landmarks in wayfinding: a review of the existing literature. Cognitive Processing 22(3):369–410 (2021).

33. Donges E, A two-level model of driver steering behavior. Human Factors 20(6):691–707 (1978).

34. Land M, Horwood J, Which parts of the road guide steering. Nature 377(6547):339–340 (1995).

35. Mars F, Driving around bends with manipulated eye-steering coordination. Journal of Vision 8(11):10–10 (2008).

36. Li WO, Saunders JA, Li L, Recruitment of a novel cue for active control depends on control dynamics. Journal of Vision 9(10):9.1–911 (2009).

37. Cruse H, Wehner R, No need for a cognitive map: decentralized memory for insect navigation. PLoS Comput Biol 7(3):e1002009 (2011).

38. Foo P, Warren WH, Duchon A, Tarr MJ, Do Humans Integrate Routes Into a Cognitive Map? Map-Versus Landmark-Based Navigation of Novel Shortcuts. Journal of Experimental Psychology: Learning, Memory, and Cognition 31(2):195–215 (2005).

39. O’keefe J, Nadel L The Hippocampus as a Cognitive Map (Oxford University Press, Oxford, 1978).

40. Tolman EC, Cognitive maps in rats and men. Psychological Review 55(4):189–208 (1948).

41. May, AJ, Ross, T. Presence and quality of navigational landmarks: effect on driver performance and implications for design. Human Factors 46(2): 346–361 (2006).

42. Simonet M et al., Probing mental representations of space through sketch mapping: a scoping review. Cognitive Research: Principles and Implications 10(1):59 (2025).

