## Supplementary material for "Learned landmark associations support online visual control under degraded visibility": SI Appendix

Grace C. Roessling
Brett R. Fajen

*Corresponding authors: Grace C. Roessling, Brett R. Fajen

### 1. Methods

**A. Dynamic Time Warping (DTW) analysis.** For each subject, two-dimensional trajectories (x- and z-coordinates) were extracted for each of the eight curved track segments in Trial 10 and Post-test 1. To compute geometric similarity, a cumulative cost matrix was constructed, representing the minimum cumulative cost of aligning every pair of points across the two trajectories. Each cell was computed iteratively from the start of both trajectories by adding the pairwise distance between points to the minimum cost of the three possible predecessor cells (corresponding to insertion, deletion, or match). The final cell, corresponding to the last points of both trajectories, contained the optimal cumulative alignment cost and thus defined the DTW distance between the trajectories. DTW cost values are bounded at zero and thus tend to be positively skewed, with a concentration of values close to zero. To address this skew, DTW costs were log-transformed prior to analysis.

**B. Rating system for drawings.** Raters were instructed to focus on five of the curved segments in the ground truth track: the chicane, the triple-S, the traffic circle, the T-turn, and the spiral. The three parabolic turns were excluded, as those turns were less geometrically distinctive and thus more difficult to identify in the drawings. The raters evaluated each drawing based on four criteria: (1) number of segments (i.e., how many segments in the drawing resemble those in the ground truth), (2) geometry (i.e., how similar in curvature and proportion are the segments present in the drawing to the ground truth), (3) order of segments (i.e., are the segments in the drawing presented in the same order as those in the ground truth); and (4) orientation (are the allocentric orientations of segments in the drawing similar to those in ground truth).

**C. Calculation of steering performance measures.** The speed of the vehicle was calculated within the Unity simulation for each frame and recorded in the log files. We used the following procedure to calculate the absolute angular steering wheel acceleration on each frame: (1) Map the output from the steering wheel device (a decimal value from -1 to +1) to the angular position of the steering wheel, which ranged from -450° to +450°. (2) Smooth the angular position data using a 10-frame rolling mean (approximately 166.7 ms). (3) Divide the difference in steering wheel angle on successive frames by the difference in time, yielding a time series of steering wheel angular velocity. (4) Smooth the angular velocity data using a 10-frame rolling mean. (5) Divide the difference in angular velocity by the difference in time on successive frames, yielding a time series of steering wheel angular acceleration. (6) Apply a 10-frame rolling mean to smooth the angular acceleration time series. (7) Take the absolute value of the angular acceleration at each frame. After each step that involved smoothing, we visually inspected the output to verify that noise was adequately removed while preserving the underlying structure of the steering wheel data.

Lane deviation was measured by calculating the vehicle's position relative to the nearest point along the road's centerline—defined as the double yellow line dividing opposing traffic lanes—and computing the signed distance between those two points. Positive lane deviation values indicated that the vehicle was to the right of the road centerline, while negative values indicate that the vehicle was to the left of the road centerline.

After calculating speed, absolute angular steering wheel acceleration, and lane deviation for each frame of each trial, we parsed each time series into sixteen segments, each of which corresponded to the track segment in which the subject was positioned for that specific frame. (Each track comprised eight straight segments interleaved with eight curved segments.) We then calculated the mean speed, standard deviation of speed, mean absolute steering wheel acceleration, and standard deviation of lane deviation for each curved segment. We excluded data from the straight track segments since our primary interest was on steering performance when the road is curved. Lastly, we averaged each metric across segments from the same visibility condition (Low or High).

2. **Figures**

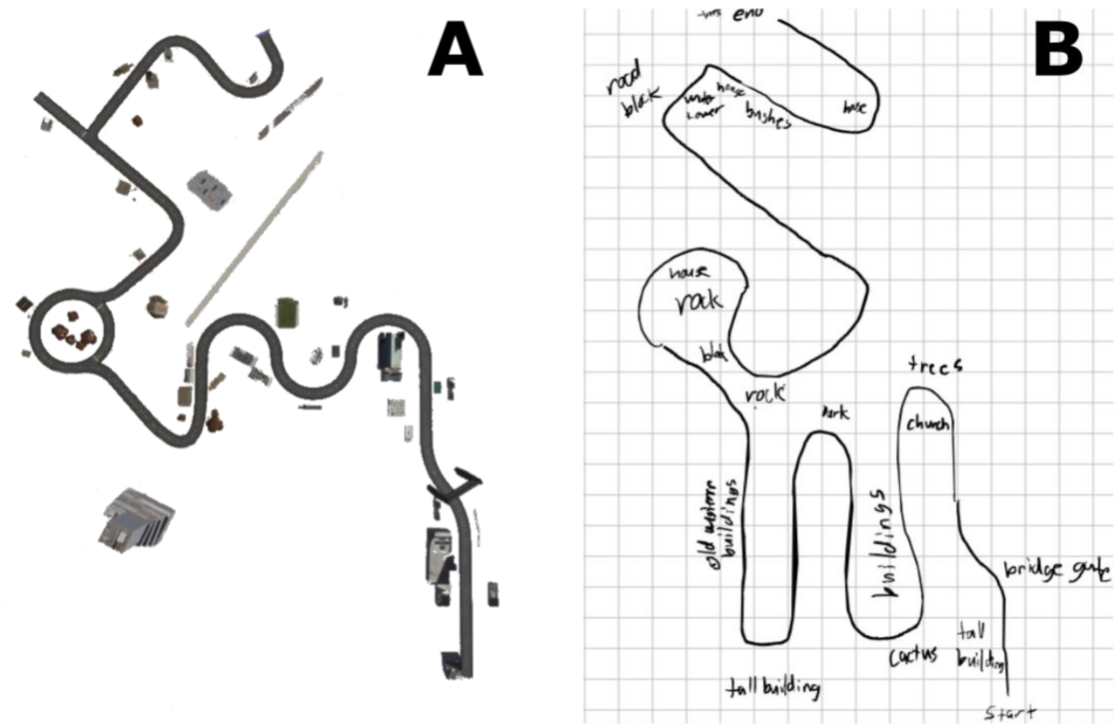


**Figure S1.** (A) Aerial view of the track in Experiment 1, along with surrounding landmarks. (B) An example of a drawing that a subject drew in Post-test 2 based on their memory of the track.

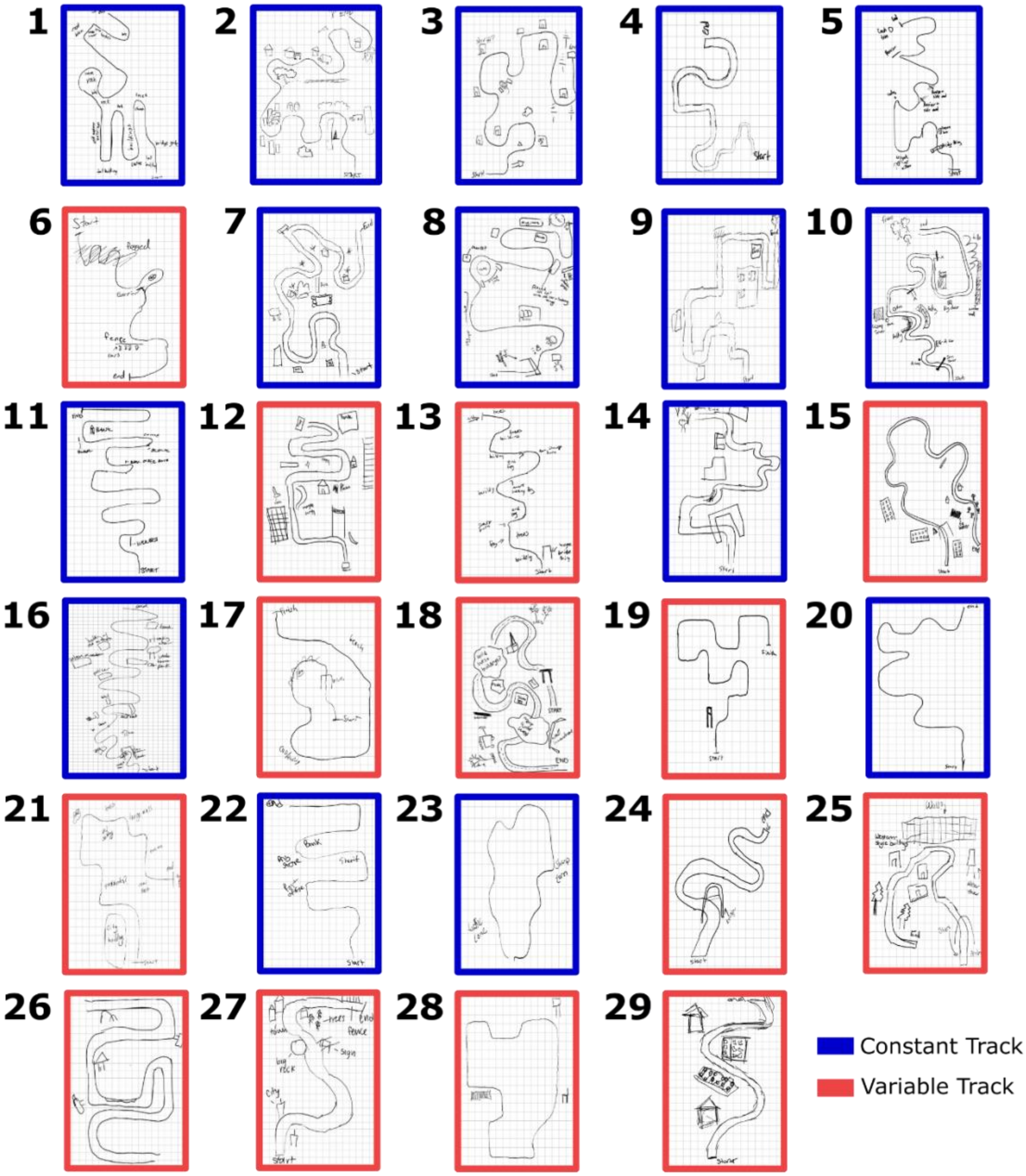


**Fig. S2.** Drawings from Post-test 2 in Experiment 1, ranked from highest to lowest score. Blue-outlined drawings are from the Constant Track group, and red-outlined drawings are from the Variable Track group.

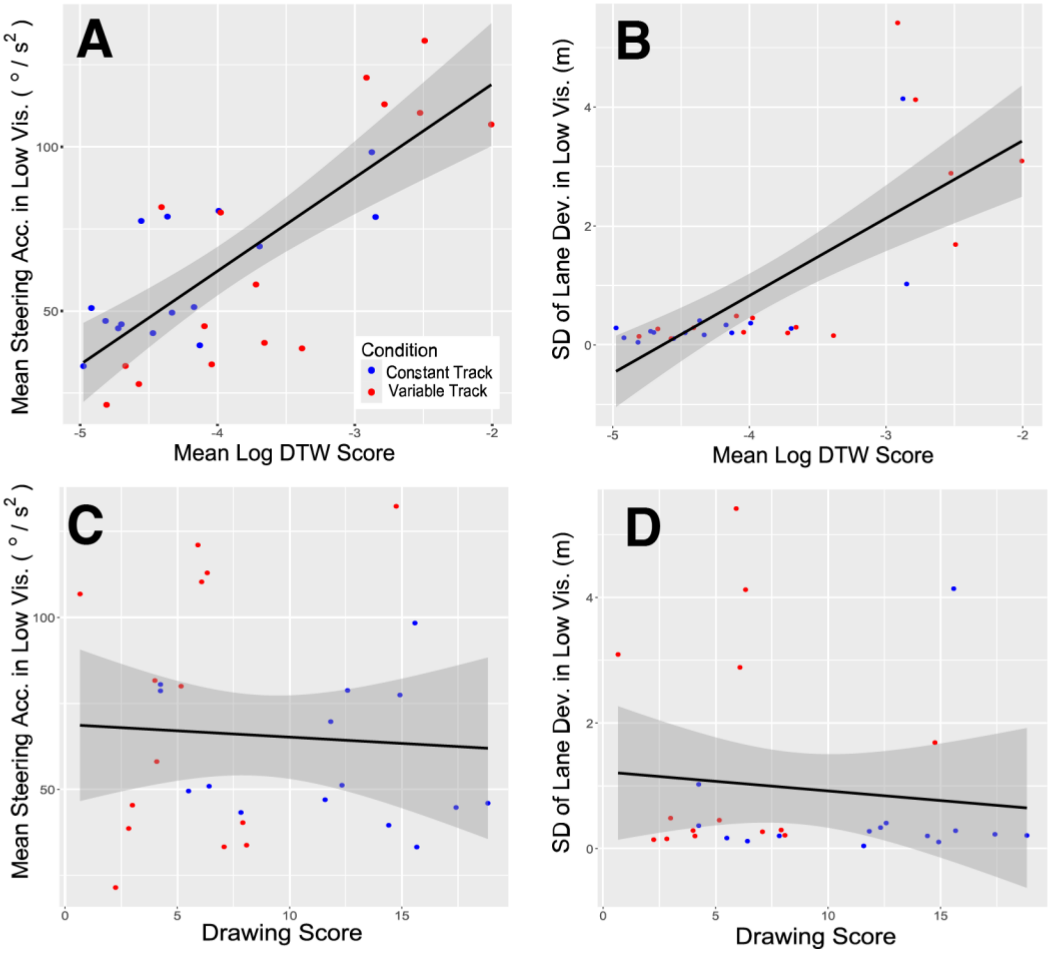


**Figure S3.** Scatterplots showing relationship between steering performance and performance on Post-test 1 and 2. Steering performance is measured in terms of mean absolute steering acceleration (A and C) and standard deviation of lane position (B and D) for low-visibility road segments on Trial 10. Performance on Post-test 1 (A and B) is measured in terms of mean log DTW score and Post-test 2 (C and D) is measured in terms of normalized drawing score.

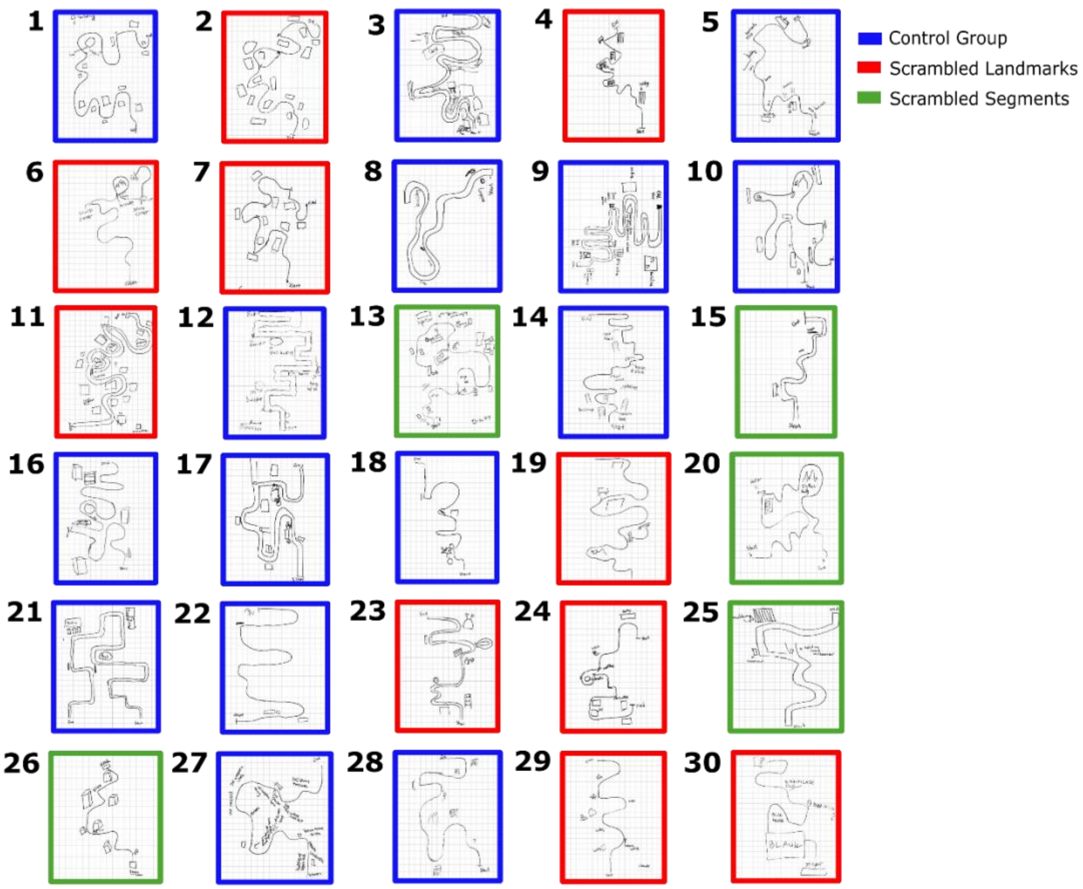


**Fig. S4.** The top 30 Post-test 2 drawings from Experiment 2, ranked from highest to lowest. Blue-outlined drawings are from the Control group, red-outlined drawings are from the Scrambled Landmarks group, and green-outlined drawings are from the Scrambled Segments group.

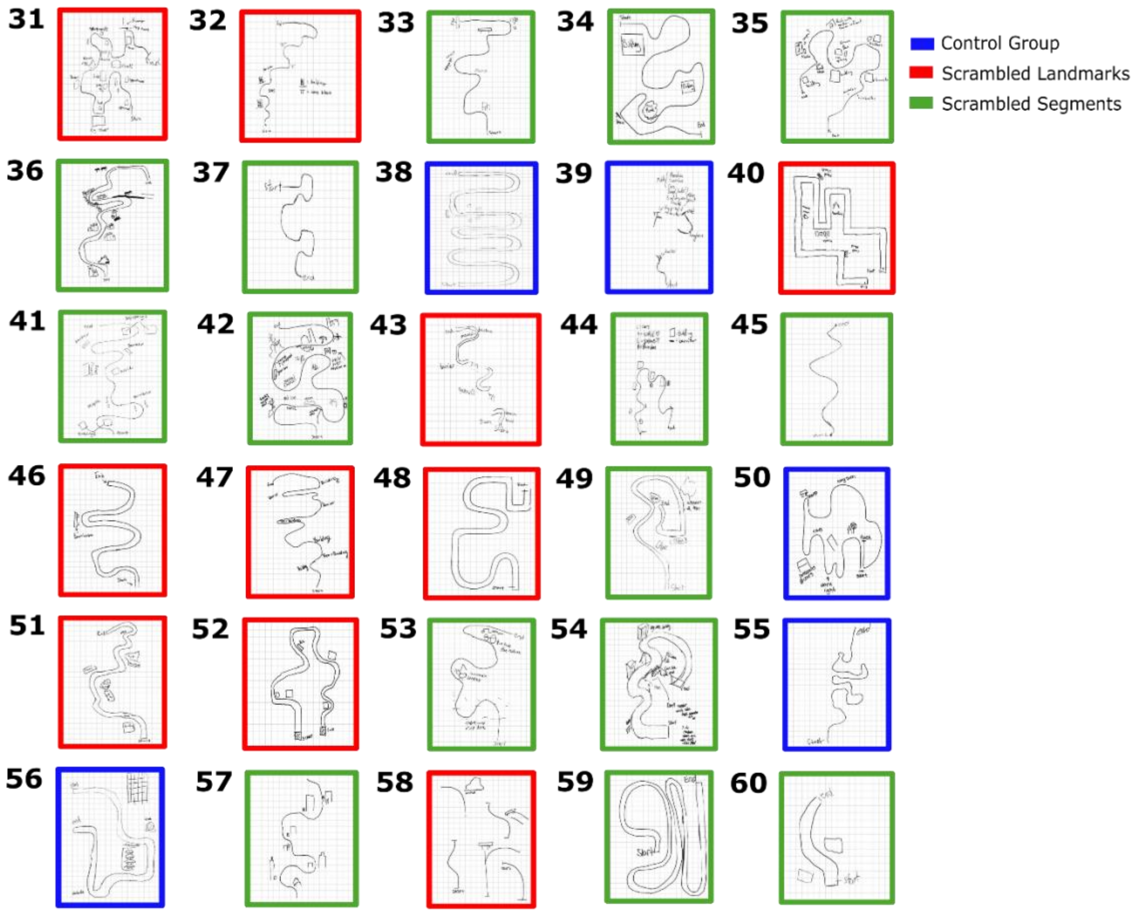


**Fig. S5.** The bottom 30 Post-test 2 drawings from Experiment 2, ranked from highest to lowest. Blue-outlined drawings are from the Control group, red-outlined drawings are from the Scrambled Landmarks group, and green-outlined drawings are from the Scrambled Segments group.

**Tables**

**Table S1.** Summary of self-reported driving and video game experience in Experiment 1.

| Category | Response | # of Subjects |
|---|---|---|
| Handedness | Right-Handed | 27 |
| | Left-Handed | 3 |
| Driver's License | Has driver's license | 27 |
| | No driver's license | 3 |
| Driving Experience | No experience | 2 |
| | Less than 1 year | 1 |
| | 1 – 2 years | 7 |
| | 2 – 5 years | 14 |
| | Over 5 years | 6 |
| Video Game Play (any genre) | Never or rarely | 9 |
| | Less than 2 hours per week | 9 |
| | 2 – 5 hours per week | 7 |
| | More than 5 hours per week | 5 |
| FPV Game Play (3D navigation) | Never or rarely | 11 |
| | Less than 2 hours per week | 12 |
| | 2 – 5 hours per week | 6 |
| | More than 5 hours per week | 1 |

**Table S2.** Summary of self-reported driving and video game experience in Experiment 2.

| Category | Response | # of Subjects |
|---|---|---|
| Handedness | Right-Handed | 53 |
| | Left-Handed | 7 |
| Driver's License | Has driver's license | 54 |
| | No driver's license | 6 |
| Driving Experience | No experience | 1 |
| | Less than 1 year | 10 |
| | 1 – 2 years | 11 |
| | 2 – 5 years | 30 |
| | Over 5 years | 8 |
| Video Game Play (any genre) | Never or rarely | 15 |
| | Less than 2 hours per week | 12 |
| | 2 – 5 hours per week | 17 |
| | More than 5 hours per week | 16 |
| FPV Game Play (3D navigation) | Never or rarely | 26 |
| | Less than 2 hours per week | 18 |
| | 2 – 5 hours per week | 9 |
| | More than 5 hours per week | 7 |

**Movie S1 (separate file).** A video showing the driver's perspective at the start of Trial 10 in Experiment 1.

**Movie S2 (separate file).** A video showing the driver's perspective at the start of Post-test 1 in Experiment 1.
